# DDK inhibition disrupts replication leading to mitotic catastrophe in Ewing sarcoma

**DOI:** 10.1101/2021.11.02.466939

**Authors:** Jeffrey C. Martin, Ajay Gupta, Tamara J. Hagoel, Lingui Gao, Miranda L. Lynch, Anna Woloszynska, Thomas Melendy, Jeremy Kane, Joseph Kuechle, Joyce E. Ohm

## Abstract

Ewing sarcoma is the second most common bone malignancy in children and adolescents. Patients with upfront metastatic or recurrent disease have poor outcomes with 5-year survival rates of <30%. CDC7, also known as DDK (DBF4-dependent kinase), is a serine-threonine kinase that, in coordination with its activation subunit ASK (or DBF4), is involved in a diverse array of cellular functions including the regulation of DNA replication initiation and activation of the replication stress response. Due to DDK’s diverse roles during replication, coupled with an increased level of genomic instability and R-loop-mediated replication stress within Ewing sarcoma cells, we hypothesized that Ewing sarcoma cells would be particularly vulnerable to DDK inhibitors. Here, we show that treatment with two selective DDK inhibitors, TAK-931 and XL413, results in apoptosis and a significant reduction in cell viability in EWS-FLI1-harboring Ewing sarcoma cell lines. We show that low dose DDK inhibition in Ewing sarcoma cells causes an accumulation of cells in late-S phase with a reduced replication capacity. There is also evidence of premature mitotic entry indicating an inability to properly complete DNA replication in a timely manner upon DDK inhibition. Also, there is a significant increase in the formation of micronuclei and other aberrant mitotic structures upon DDK inhibition in Ewing sarcoma cells indicating a failure to properly progress through S-phase followed by improper mitotic entry/progression, resulting in mitotic catastrophe. Interestingly, we observed minimal signs of mitotic accumulation, despite clear evidence of replication and mitotic stress, suggesting a failure to properly enforce the mitotic checkpoint. Together, these results suggest that Ewing sarcoma cells rely on the activity of DDK to maintain cell viability and suggest that DDK inhibition may prove to be a viable therapeutic strategy for patients with Ewing sarcoma.

## Introduction

Ewing sarcoma is the second most common bone malignancy with a peak incidence in adolescent and young adult (AYA) patients ^1^. For patients that present with metastatic or recurrent disease, the 5-year survival rate is less than 30% ^2^. Upon diagnosis, the standard of care approach for the treatment of Ewing sarcoma (irrespective of localized vs metastatic disease) is neoadjuvant chemotherapy, followed by local control (definitive surgery +/- radiotherapy), then consolidation with adjuvant chemotherapy ^2^. Chemotherapy regimens vary. In North America, standard Ewing chemotherapy consists of 5-drug therapy (VDC-IE: Vincristine/Doxorubicin/Cyclophosphamide alternating with cycles of IE or Ifosfamide/Etoposide). Much work has been done to understand the molecular mechanisms that lead to Ewing sarcoma, but little headway has been made towards the development of effective targeted therapies ^3,4^. With this reality, standard of care treatment is often accompanied by debilitating side effects that can dramatically impact the patient’s quality of life ^1^. Considering the prevalence of Ewing sarcoma in the AYA population, it is important to minimize exposure to cytotoxic chemotherapy agents (e.g., alkylating agents or etoposide) that may increase the risk for secondary malignancies. Unfortunately, ∼10% of Ewing sarcoma survivors will develop treatment-associated secondary cancers at some point in their life, and a significant proportion will develop long-term complications such as cardiomyopathy and infertility ^5-7^. It is important that studies be conducted to elucidate the etiology of Ewing sarcoma and molecular mechanisms driving this cancer, as these may lead to the development of novel, less toxic therapeutic options.

The term replication stress (RS) refers to molecular processes that obstruct the progression of the replisome during replication ^8^. RS increases the likelihood of errors during replication and therefore poses a direct threat to the fidelity of the genetic material. RS is known to accelerate the oncogenic process and is a frequent outcome of aberrant oncogene activity ^8^. Examples of known molecular events impeding replication include things such as RNA:DNA hybrids (R-loops) formed during transcription ^9^, protein-DNA barriers that impede the progression of the helicase/polymerase ^10^ or exhaustion of nucleotides caused by a deregulation in nucleotide synthesis/consumption ^11^. Each of these processes form barriers, physical and/or functional, to the processivity of the replicative helicase and can lead to prolonged fork stalling and fork collapse resulting in the formation of DNA double strand breaks. Fork stalling must be accompanied by proper regulation of the replisome components in order to enable efficient restart and completion of genome duplication ^8^. The intra-S phase checkpoint, mediated by the Ataxia telangiectasia and Rad3-related (ATR) kinase and its effector kinase CHK1 is the main mechanism that the cell utilizes to deal with this type of stress ^12^. Exacerbating the level of RS through the inhibition of the RS response or treatment with replication inhibiting agents such as aphidicolin or hydroxyurea has proven to be a very effective treatment strategy for some tumor types ^13^.

There is a growing body of evidence implicating RS as a major source of genomic instability in Ewing sarcoma ^14^. For example, Ewing sarcoma cell lines and xenografts have been shown to be sensitive to ATR inhibitors as single agents ^15^ and in combination with the inhibition of WEE1 ^16^, suggesting a greater dependency on these enzymes. Also, the inhibition of nucleotide synthesis with the use of hydroxyurea or gemcitabine results in DNA damage and cell death at very low concentrations, and these compounds synergize with inhibitors of CHK1 ^17^ suggesting a deficiency in fork stability and/or restart. Importantly, it was recently shown that an EWS-FLI1-mediated increase in CDK9 activity and RNAPII transcription results in the formation of R-loops and blocks BRCA1-mediated homologous recombination repair ^18^. R-loop formation poses a serious threat to the processivity of the replisome and is a major source of RS in certain tumors ^19^. Elimination of genomic R-loops by overexpression of RNaseH1 desensitized Ewing sarcoma cells to the inhibition of ATR ^18^, suggesting that R-loop formation plays a large role in the accumulation of RS within these cells. Due to the intrinsic level of RS within these cells, we reasoned that it is likely that targeting of the RS response may prove to be a viable treatment strategy for Ewing sarcoma.

CDC7 (cell division cycle 7) is a serine-threonine kinase that, together with its activation subunit ASK (or DBF4), is involved in a diverse array of cellular functions. CDC7 is commonly referred to as DDK (DBF4-dependent kinase) and its primary role is in the regulation of the initiation of DNA replication ^20^ through the direct phosphorylation of several members of the MCM helicase complex ^21,22^. It has classically been thought to be inactivated during times of RS ^23,24^. This is believed to be in an attempt to reduce the level of active replication forks, limiting the capacity of cells to accumulate further RS. However, several recent publications have highlighted an active role for DDK during RS. For example, it is required for the full activation of CHK1 through its phosphorylation of the adaptor protein CLASPIN ^25^. DDK also appears to play a crucial role at stalled replication forks ^26^ as several of its phosphorylation targets have been implicated in alternative fork stability/restart mechanisms ^27-29^. It is also important to note that the activation of dormant replication origins in the vicinity of stalled forks to rescue replication of the local DNA would require DDK activity. It is, therefore, likely that apart from its active role in the RS stress response activation, DDK also plays a somewhat passive role in the protection of stalled forks through the activation of adjacent dormant origins.

Due to the intrinsic level of RS within Ewing cells, and the central role of DDK in the RS response, we hypothesized that Ewing cells would be reliant on DDK for proper replication and cell division. Here we provide evidence that DDK inhibition within Ewing cells results in failure to properly complete DNA replication, mitotic catastrophe, and apoptotic cell death. This study reveals the potential to inhibit DDK for the treatment of Ewing sarcoma and provides further evidence that these tumors are sensitive to the inhibition of the RS response.

## Results

Due to the protective role that DDK plays against RS, we hypothesized that Ewing sarcoma cells would be particularly sensitive to the inhibition of DDK. To test this, we treated three Ewing sarcoma cell lines (A673, RD-ES, and SK-ES-1) with increasing concentrations of two independent DDK inhibitors, XL413 and TAK-931, for 72 hours and assessed cell viability. TAK-931 has been shown to be a much more specific inhibitor of DDK ^30^ and was recently tested in a phase 2 clinical trial for the treatment of metastatic pancreatic cancer, metastatic colorectal cancer and advanced solid tumors. The osteosarcoma cell line, U2OS, was used as a non-Ewing sarcoma/bone tumor control in all the following studies.

We found that Ewing sarcoma lines were significantly more sensitive to both DDK inhibitors as compared to the U2OS osteosarcoma cell line (**Figure 1A**). Using an initial range of concentrations of XL413 between 1 and 10 µM ^30^, we did not see any statistically significant reduction in U2OS viability at even the highest concentrations. In the Ewing sarcoma lines, cell viability was reduced in a statistically significant, dose dependent manner. A significant reduction in cell viability was observed even at the lowest concentration tested (1µM). Viability at 72 hours was reduced to 68.2% in A673 (p=0.0002), 72.6% in RD-ES (p=0.001), and 54.1% in SK-ES-1 (p<0.0001) with 1 µM XL413 (Figure 1A, left panel). Similar results were obtained with TAK-931 (Figure 1A, right panel). We used a range between 300nM and 5µM TAK-931, and while the U2OS cells do respond to higher concentrations of this compound, the Ewing sarcoma cell lines remain remarkably more responsive. At the lowest dose tested (300nM), the U2OS line shows a viability of 90.8%, while the Ewing sarcoma lines are reduced to 18.3% (p<0.0001), 29.5% (p<0.0001) and 30.1% (p<0.0001) in A673, RD-ES, and SK-ES-1, respectively. To ensure this was a phenomenon shared among several Ewing sarcoma cell lines, a wider range of Ewing sarcoma cell lines with varying genetic backgrounds (TP53 and STAG2 WT vs. mutant) were tested. Cells were treated with TAK-931 for 72 hours and IC50 values were calculated (**supplemental figure 1A**). In our hands, all Ewing sarcoma cell lines tested had IC50s below that of U2OS (5.33µM) (**supplemental figure 1B & C**). Importantly, there was no apparent relationship between TP53, STAG2 or CDKN2A status and response to TAK-931, suggesting that the basis for this sensitivity is independent of the function of these proteins (**supplemental figure 1C, D & E**).

**Figure 1.**
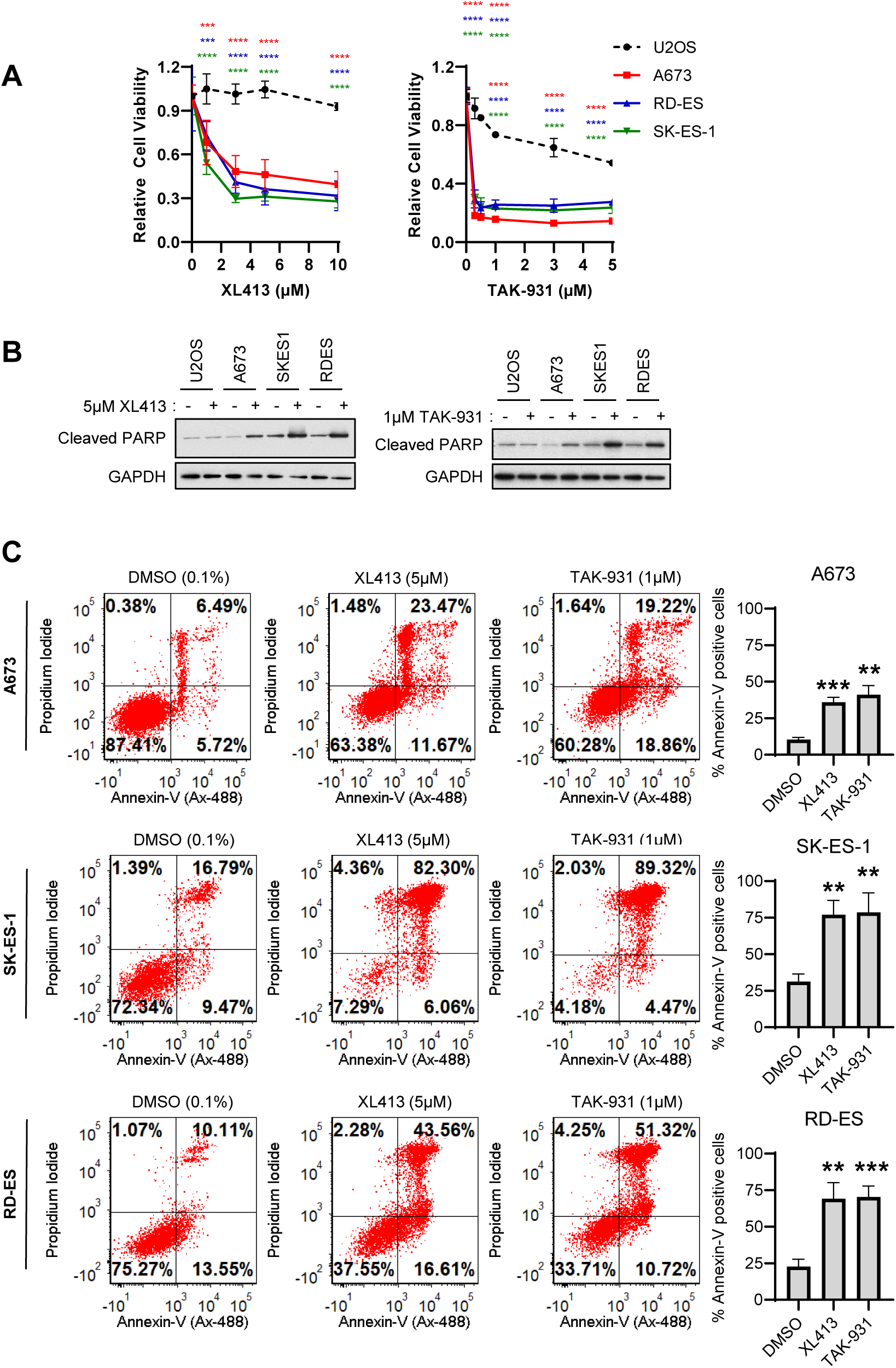
Ewing Sarcoma cells are sensitive to DDK inhibition. A) Ewing Sarcoma cells (A673, RD-ES and SK-ES-1) and the non-Ewing, osteosarcoma cell line U2OS were treated with increasing concentrations of the DDK inhibitors XL413 and TAK-931 for 72 hours and cell viability was measured using CCK-8 cell viability reagent. Relative viability was calculated based on DMSO treated cells (n=3 technical replicates). 2-way ANOVA multiple comparisons, ***p<0.001, ****p<0.0001.B) Cells were treated with either 0.1% DMSO, 5µM XL413 (left panel) or 1µM TAK-931 (right panel) for 48 hours. Whole cell lysates were collected, and western blot was performed for the specified proteins. GAPDH was used as a loading control (representative image of 3 independent experiments).C) Ewing Sarcoma cells were treated with 0.1% DMSO, 5µM XL413 or 1µM TAK-931 for 48 hours. Cells were stained with anti-Annexin-V (AlexaFluor-488 conjugated) and propidium iodide and analyzed using flow cytometry. Apoptotic cells were designated as Annexin-V (upper right and lower right quadrants) positive cells and were measured using FCS express v7 (n=3 biological replicates). Unpaired t test, ***p<0.001, **p<0.01.

To determine if there was an induction of apoptosis upon DDK inhibition, Ewing sarcoma cells were treated with either 5µM XL413 or 1µM TAK-931 for 48 hours and induction of apoptosis was then assessed by evaluating cleaved parp (indicator of caspase activity) (**Figure 1B**) and Annexin-V/PI staining (**Figure 1C**). These doses were chosen based on the cell viability experiments in figure 1 and supplemental figure 1 and specifically because these were the lowest doses at which we observed maximal reduction in cell viability in most Ewing cell lines. We found that both XL413 and TAK-931 induced apoptosis at 48 hours in the Ewing sarcoma cells (**Figure 1B & C**). We observed an increase in cleaved-PARP by western blot in all three Ewing cell lines upon treatment with XL413 (**Figure 1B, left panel**) and TAK-931 (**Figure 1B, right panel**). No similar induction of cleaved-PARP was observed in U2OS. We also performed Annexin-V/PI staining of our cell lines in response to DDK inhibitor treatment. In A673 cells compared to DMSO vehicle, XL413 increases Annexin V positive cells from 10.5% to 36% (p=0.0003) and TAK-931 increases Annexin-V positive cells to 41.2% (p=0.0012). This is also observed in RD-ES cells (increase in Annexin V positive cells from 22.7% to 69.1%; p=0.0028 with XL413 and 70.4%; p=0.0008 with TAK-931) and SK-ES-1 cells (increase in Annexin V positive cells from 31.3% to 77%; p=0.002 with XL413 and 78.58; p=.0046 with TAK-931). This suggests an active role of DDK in the prevention of apoptosis in Ewing sarcoma cells.

Due to DDK’s role in genome maintenance during replication ^31^, apoptotic cell death that occurs upon its inhibition is likely a result of an increased level of RS ^30^. We hypothesized that due to the intrinsic level of RS within Ewing sarcoma cells, they would have a substantial reliance on DDK activity for proper S-phase progression. If this hypothesis is correct, relatively low doses of DDK inhibitors would result in an apparent increase in RS and reduction in replication rates. A common output of RS is the phosphorylation of ATR substrates, most notably CHK1. However, DDK has been shown to be required for the full activation of CHK1 ^25^ and ATR ^27^, therefore, assessing ATR activity upon DDK inhibition is not a viable approach. Instead, to test whether DDK inhibition results in RS in Ewing sarcoma cells, we measured the level of incorporation of the nucleotide analog EdU after 8-hour DDK inhibition. Due to DDK’s role in the activation of replication origins, a reduction in replication rates is expected upon its inhibition. However, aberrant levels of replication stalling will lead to dramatic and non-uniform reductions in the EdU incorporation rate, even upon a short treatment pulse. As expected, in U2OS cells, both TAK-931 (300nM) and XL413 (1µM) treatment led to a small and uniform reduction in EdU incorporation as measured by flow cytometry (**Figure 2A, top panel**). However, DDK inhibition results in a dramatic and non-uniform reduction in EdU incorporation in all three Ewing sarcoma lines. In these cells, the EdU staining shifts to the left, and the peaks become broader, indicating aberrant replication slowing in a subset of cells, suggesting increased RS (**Figure 2A, lower three panels**).

**Figure 2.**
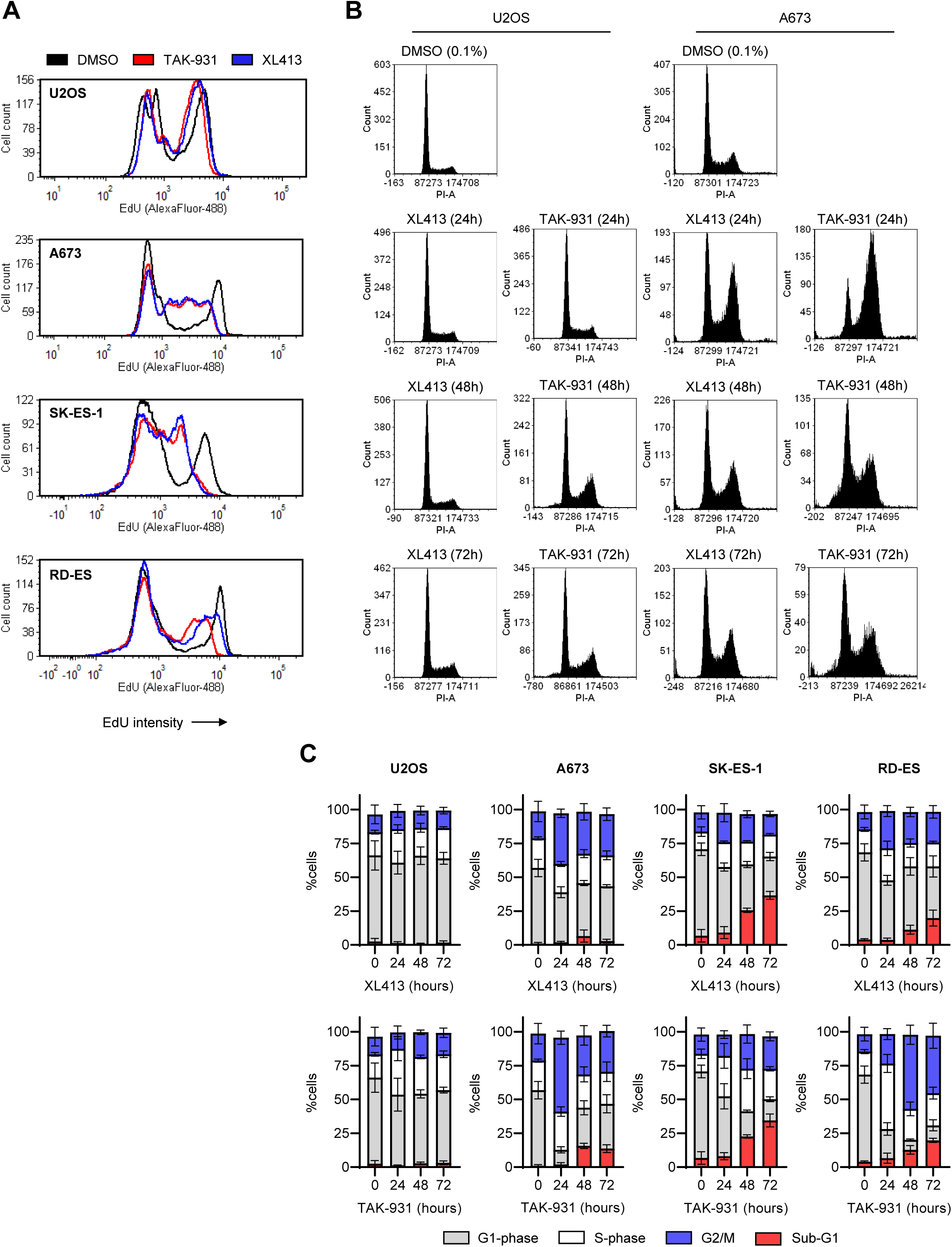
DDK inhibition induces RS and causes aberrant cell cycle accumulation in Ewing Sarcoma cells. A) Ewing Sarcoma cells and U2OS cells were treated with either 0.1% DMSO (black line), 300nM TAK-931 (red line) or 5µM XL413 (blue line) for 8 hours. Then, in the presence of DDKi, cells were incubated with 10µM EdU for 30 minutes. Cells were then fixed and stained for EdU (AlexaFluor488) (X-axis) and EdU intensity was analyzed using flow cytometry. Representative image of three independent experiments. B & C) Ewing Sarcoma cells and U2OS cells were treated with 0.1% DMSO (72 hours), 300nM TAK-931 or 1µM XL413 for 24, 48 and 72 hours, DNA was stained with propidium iodide and DNA content was analyzed using flow cytometry. Cell cycle distribution was measured using FCS Express v7. B) Representative histograms of three independent experiments showing only the results obtained with A673 and U2OS cells. C) Quantification of cell cycle distribution; G1 = grey bars, S = white bars, G2/M = blue bars, sub-G1 = red bars (n=3 biological replicates).

We were then interested in determining the effect of DDK inhibition on cell cycle progression in Ewing cells. Asynchronous A673 and U2OS cells were treated with TAK-931 (300nM) or XL413 (1µM) for 24, 48 and 72 hours and cell cycle was analyzed. The U2OS cells experienced minimal cell cycle changes upon DDK inhibition at all timepoints tested. However, at 24 hours of DDK inhibition, the Ewing sarcoma cells accumulated in late-S and G2 phases of the cell cycle (**Figure 2B & C**) which is consistent with the reduction in EdU incorporation seen in figure 2A. Interestingly, there is a reduction in G2 cells by 48 hours (% cells in G2 reduced to 28.77% at 48 hours TAK-931 and 30.95% XL413) and this is accompanied by the emergence of sub-G1 DNA content (%sub-G1 cells upon TAK-931 treatment: 0 hours = 1.0%, 24 hours = 2.01% (p=.8809), 48 hours = 15.7%(p<0.0001), 72 hours = 13.7%(p=0.0001)) (**Figure 2B & C**). This phenomenon was better represented upon TAK-931 treatment which is consistent with this compound being a more potent inhibitor of DDK. Similar results were seen in the SK-ES-1 and RD-ES cell lines (**Figure 2C & supplemental figure 3**) suggesting that this is most likely a phenomenon that is conserved across most Ewing sarcoma cell lines. Overall, these results suggest that DDK plays a large role in the conservation of DNA replication rates and S-phase progression in Ewing cells.

To gain more insight into the role of DDK in replication progression in Ewing sarcoma, A673 cells were synchronized at the G1-S transition using a double-thymidine block (DTB) and then released into S-phase (dT-release) +/- 300nM TAK-931 (**Figure 3A**). In line with the data in figure 2, DDK inhibition significantly prolonged S-phase, ∼4-8 hours longer than DMSO treated cells (**Figure 3B, 3C & 3D & 3E middle**). DMSO treated cells completed replication (peak in % cells w/ 4N DNA content) between 8-12 hours upon dT-release while TAK-931 treated cells did not complete replication until ∼20 hours upon dT-release (**Figure 3E right**). Also, TAK-931 treatment seemed to disrupt the transition of cells from S-phase into mitosis. This is evidenced by the fact that cells seemed to reside in G2/M for a prolonged period compared to the DMSO treated cells, suggesting delayed mitotic entry/progression (**Figure 3E, right panel**). Also, in the TAK-931 treated group, there appeared to be cells with sub-4N DNA content even at very late timepoints (16-20 hours post dT-release) (**Figure 3B & 3E middle panel**) suggesting a significant portion of cells may not complete DNA replication prior to mitotic entry.

**Figure 3.**
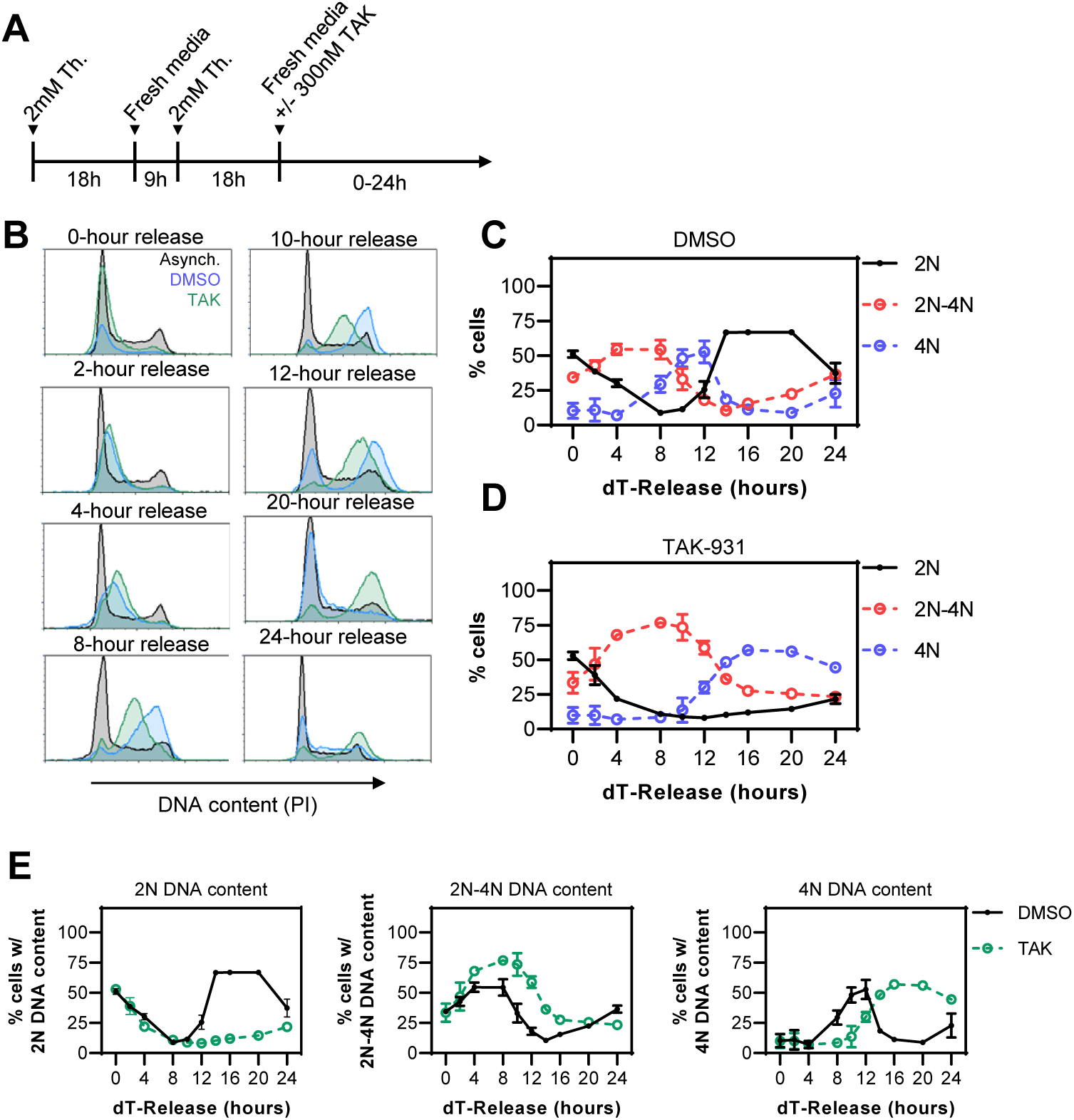
DDK inhibition results in prolonged S and G2/M phases in Ewing sarcoma cells. A) A673 cells were subjected to a double thymidine block (described in methods) and then released (dT-release) into either fresh media or media containing 300nM TAK-931 for various time points. Cell cycle phase was visualized by measuring DNA content with flow cytometry. B) Representative histograms upon dT-release at indicated timepoints. Histogram colors: grey – asynchronous; blue – 0.1% DMSO treated cells; green – 300nM TAK-931 treated cells. Representative images of 2 independent experiments. C & D) Quantification of % cells with either 2N (black), 2-4N (red) or 4N (blue) DNA content over the course of 28 hours post dT-release. DMSO treated cells (C), 300nM TAK-931 treated cells (D) (n = 2 biological replicates; 14h,16h, 20h are n=1). E) Quantification of % cells with 2N (top) 2-4N (middle) or 4N (bottom) DNA content. DMSO treated cells – black line; 300nM TAK-931 treated cells – green line (n = 2 biological replicates; 14h,16h, 20h are n=1).

To determine if DDK inhibition results in premature mitotic entry in Ewing sarcoma cells, A673 cells were treated with 300nM of the DDK inhibitor TAK-931 for 24 and 48 hours. Cells were then stained for DNA content as well as for the mitotic marker phospho-histone H3 (ser3) (p-HH3). We observed a significant increase in the proportion of p-HH3-positive cells with sub-G2 DNA content upon DDK treatment (**Figure 4A & 4B**) from 3.33% with no treatment to 16.2% at 24 hours (p=0.0094) and 17.9% at 48 hours (p=0.0051), suggesting premature mitotic entry. Interestingly, there was a slight but statistically significant reduction in total p-HH3-positive cells at 24 (43% reduction; p=0.0121) and 48 hours (43% reduction; p=0.0117) of DDK inhibition (**Figure 4C & 4D**) suggesting a lack of mitotic delay. These results were confirmed by western blot. Nocodazole treatment was used as a positive control to mark mitotic accumulation. Nocodazole induces a large increase in the level of pHH3 as is to be expected of cells enforcing a spindle-assembly checkpoint (SAC) but there is no apparent increase upon 24 or 48-hour TAK-931 treatment (**Figure 4E**) despite clear evidence of cell cycle alterations at these timepoints (**Figure 2B**). This was a puzzling find because mitotic delay is a common outcome of cells that are experiencing RS ^32^ and TAK-931 has been shown to cause mitotic arrest in other cancer cell lines ^30^.

**Figure 4.**
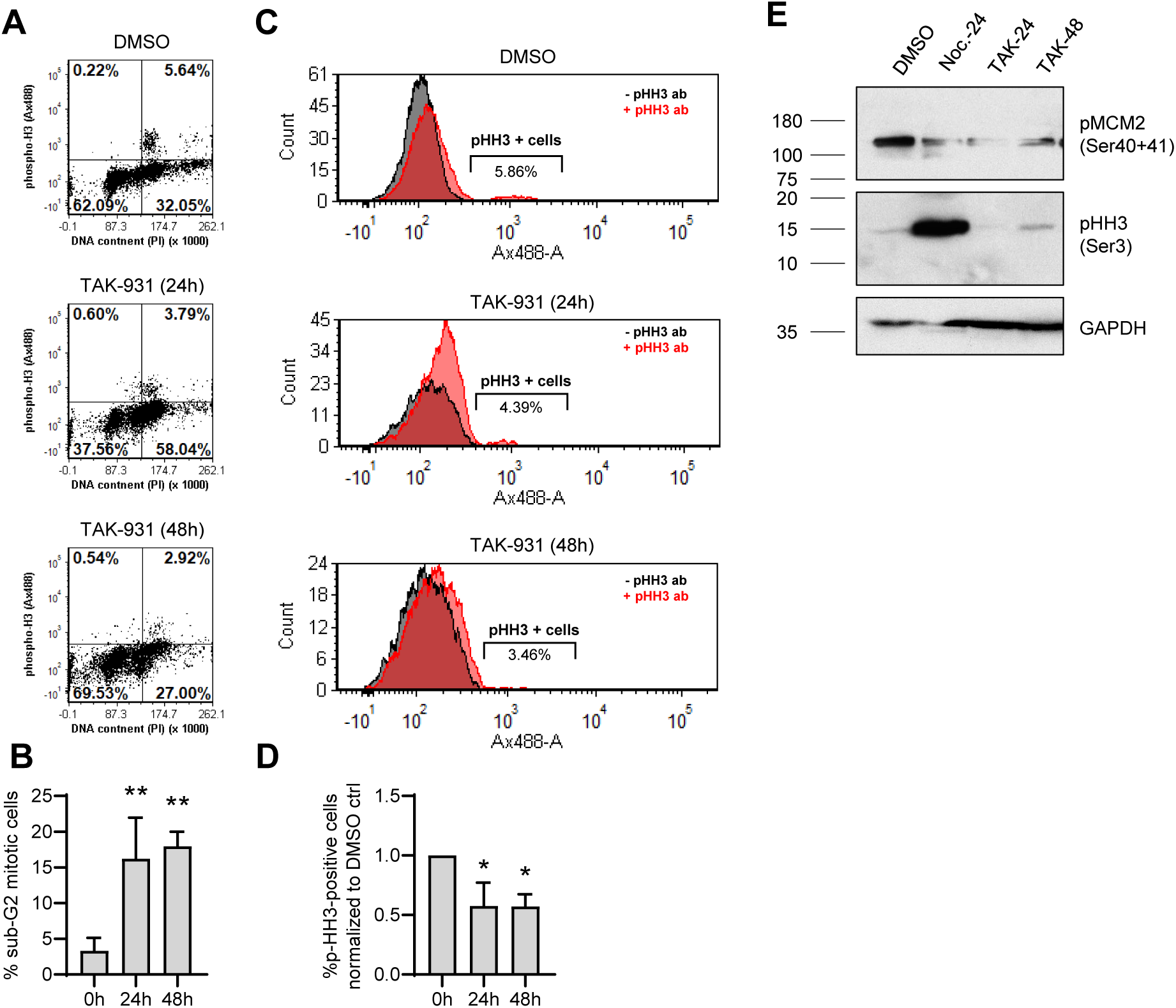
DDK inhibition results in premature mitotic entry in Ewing sarcoma cells. A) A673 cells were treated with 0.1% DMSO (48 hours) or 300nM TAK-931 for 24 and 48 hours. Cells were then fixed and stained for phospho-Histone H3 (Ser10) (AlexaFluor488) (designated as pHH3) and DNA (Propidium Iodide) and analyzed using flow cytometry. B) The percentage of sub-G2 mitotic cells was calculated by dividing the number of sub-G2 cells that stained positive for pHH3 (top left quadrant of dot plot) by the total number of pHH3-positive cells and multiplying by 100 (n=3 biological replicates). Ordinary one-way ANOVA compared to DMSO treated control, **p<0.001. C & D) The percentage of pHH3-positive cells was identified as cells that showed positive staining with the addition of pHH3 antibody (red histogram) that did not stain positive without the addition of a pHH3 antibody (black histogram) (n=3 biological replicates). Ordinary one-way ANOVA compared to DMSO treated control, *p<0.05. E) A673 cells were treated with 1µM Nocodazole for 24 hours or 300nM TAK-931 for 24 or 48 hours. Protein lysates were collected, and a western blot was performed for specified protein targets. GAPDH was used as a loading control (Western blot validation, n=1.)

Under-replicated DNA entering mitosis due to RS or failure to properly enforce SAC would result in mitotic catastrophe ^33^. This can be observed through the appearance of abnormal mitotic structures such as anaphase bridges, lagging chromosomes and the disorganization of centrosomes due to the uncoupling of DNA replication and mitosis ^30^. To this end, we observed a marked increase in the number of abnormal mitotic structures upon DDK inhibition in A673 cells (**Figure 5A**). These structures were not observed in basal conditions and seemed to appear most dramatically at 48 hours of treatment. These include cells with anaphase bridges (**Figure 5A.a**), lagging chromosomes (**Figure 5A.b**), cells with >2 centrosomes (**Figure 5A.c and 5A.d**) and other unclassified abnormal mitotic structures (**Figure 5A.e & 5A.f**). We also observed a significant increase in the formation of micronuclei upon DDK inhibition (**Figure 5B**). Micronuclei form in daughter cells due to improper chromosomal separation and are indicative of mitotic catastrophe ^34^. Consistent with the results seen in figure 2, the appearance of micronuclei did not occur until 48 hours of treatment (increase from 4.7% to 23.1% (p<0.0001) and 15.45% (p=0.0025) with TAK-931 and XL413 respectively), suggesting that, despite the initial S-phase accumulation, cells proceed through the cell cycle as planned and attempt to undergo cell division. Mitotic catastrophe results in the accumulation of DNA damage in the subsequent G1 phase ^35^. To this end, there was a marked increase in the level of gamma-H2AX staining (indicative of DNA double strand breaks) at 24 hours (p=0.0153) 48 hours (p<0.0001) and 72 hours (p<0.0001) of TAK-931 treatment and 48 hours (p<0.0001) and 72 hours (p<0.0001) of XL413 treatment (**Figure 5C & 5D**). Importantly, the most significant induction of damage occurred at later time points, consistent with the idea that the induction of DNA damage requires a round of cell division. Unexpectedly, DDK inhibition did not result in the generation of aneuploid (>4N DNA content) cells (**supplemental figure 6**). This may be due to the inability of the cells that acquired additional genetic material to complete the subsequent S-phase before undergoing apoptosis. Nonetheless, these results support the idea that, upon DDK inhibition, Ewing sarcoma cells undergo mitotic catastrophe and cell death.

**Figure 5.**
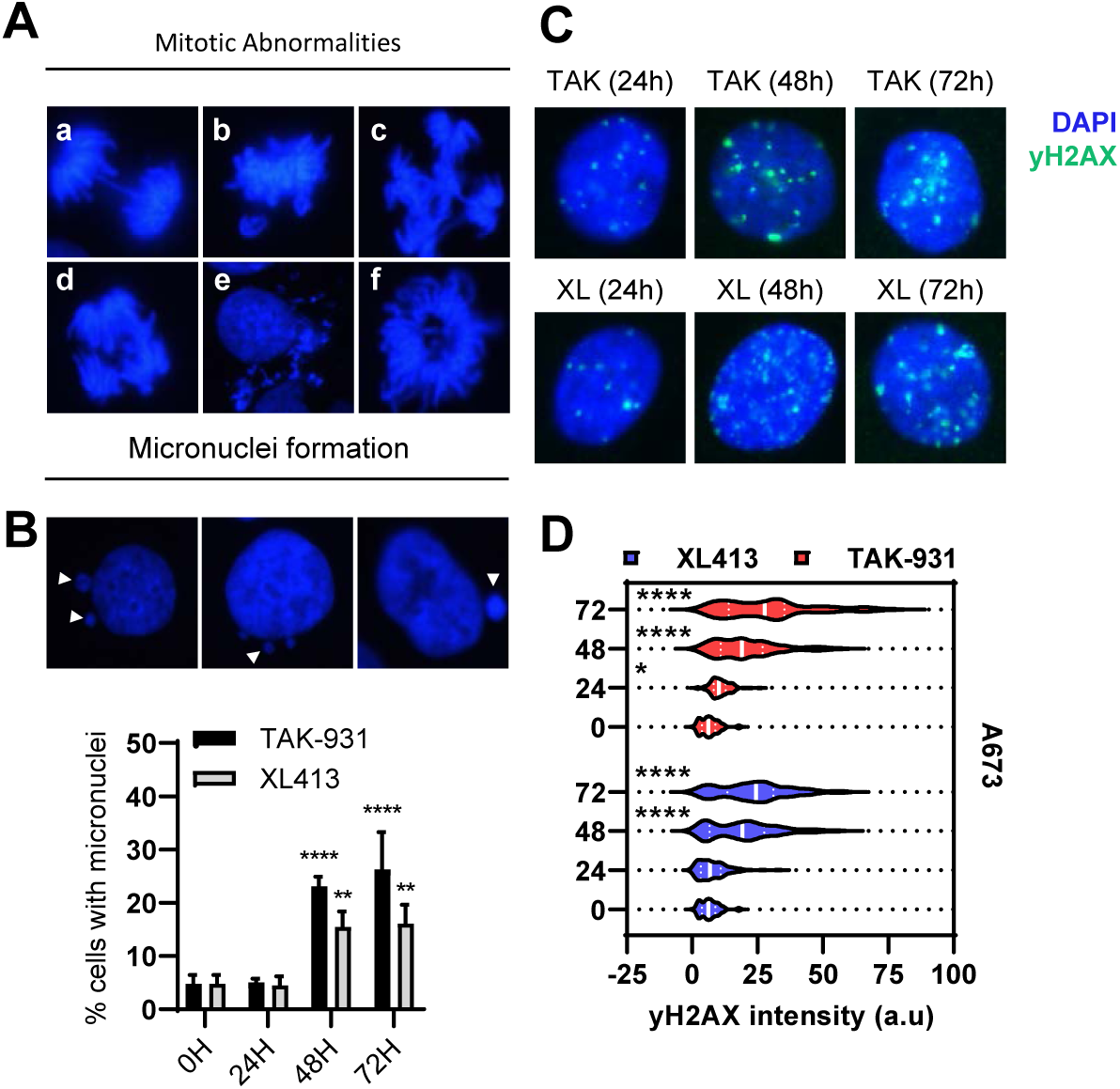
DDK inhibition causes mitotic defects and DNA damage in Ewing sarcoma cells. A) Representative images of mitotic abnormalities induced in A673 cells upon DDKi treatment. a) anaphase bridges, b) lagging chromosomes, c) four centrosomes, d) three centrosomes, e) mitotic catastrophe, f) unclassified mitotic abnormality (representative images of two independent experiments). B) A673 cells were treated with 0.1% DMSO (72 hours), 300nM TAK-931 or 1µM XL413 for 24, 48 and 72 hours and DNA content was stained with DAPI. Percent cells with micronuclei were quantified as any cell that displayed 1 or more micronuclei (white arrows) structure around the nucleus (n = 3 biological replicates). 2way ANOVA, ****p<0.0001, **p<0.01. C) A673 cells were treated with either 0.1% DMSO, 300nM TAK-931 or 1µM XL413 for 24, 48 and 72 hours. Cells were fixed and stained for yH2AX (AlexaFluor 488 – green) and DNA (DAPI – blue). D) Corrected total cellular fluorescence intensity for yH2AX was calculated using FIJI image analysis software (representative images of two biological replicates ∼100 cells were quantified per condition). 2way ANOVA, *p<0.05, ****p<0.0001.

## Discussion

In this study, we show that two independent inhibitors of DDK (XL413 and TAK-931) significantly reduce proliferation and cause cell death by apoptosis in Ewing sarcoma cells. We also show that DDK inhibition resulted in a non-uniform reduction in replication rates in Ewing sarcoma cells indicative of RS. This results in an accumulation of cells in the S and G2-phases of the cell cycle after 24 hours of treatment. Furthermore, DDK inhibition leads to the accumulation of mitotic abnormalities such as anaphase bridges and multi-poled cells at 48 hours of treatment, most likely due to under-replicated DNA entering mitosis. There was also a large increase in the number of cells with micronuclei formation, indicative of mitotic catastrophe. Interestingly, DDK inhibition did not result in an increase in the number of cells in mitosis despite clear evidence of replication and mitotic stress, suggesting a failure to properly enforce a mitotic checkpoint. While the sensitivity too DDK inhibition is clear, additional work is needed to understand DDK’s specific role in replication progression in Ewing sarcoma.

One potential mechanism is that DDK inhibition disrupts the activation of dormant origins in response to elevated levels of RS in Ewing sarcoma. Dormant origins play an important role in the prevention of replication stress-induced genomic instability ^36^. Resolution of a stalled replication fork requires the concerted action of a growing number of proteins ^8^. However, a stalled replication fork need not be resolved if an adjacent fork is active and can complete replication of the vicinal DNA. However, upon the occurrence of two adjacent stalled replication forks, the genomic region that resides between them may fail to be replicated. In this scenario, proper mitosis is not possible as the sister chromatids will not fully separate. Mitotic progression under these conditions will most likely result in the formation of anaphase bridges and other abnormal mitotic structures ^37^. While this may be the underlying source of DDKi sensitivity in Ewing sarcoma, it is unlikely the case. This is because DDK and CDK2 work in conjunction to activate replication origins. To this point, Ewing sarcoma cells lines were not significantly sensitive to the CDK2 inhibitor Roscovitine and high dose Roscovitine treatment did not significantly alter cell cycle progression in any way (**supplemental figure 4**). However, it is possible that CDK2 and DDK play disproportionate roles in the activation of dormant origins, with DDK carrying a heavier load in this regard. It is also important to point out that the molecular events that lead to the activation of dormant origins are likely separate from those that act at scheduled origins, as dormant origins are typically fired during times of replication stress when global replication progression is halted ^38^. Interestingly, the firing of dormant origins has been linked to CHK1 activity and, as mentioned above, DDK plays a large role in the activation of CHK1 ^25^. Therefore, it remains possible that DDK plays a larger role in the specific activation of dormant origins than CDK2 which may explain why Ewing cells do not rely on CDK2 to such an extent.

While the potential to utilize dormant origins provides a safeguard against RS-induced genomic instability, fork stability and restart is arguably the most fundamental. Apart from the activation of replication origins, recent work has highlighted a role for DDK in the activation of nucleases involved in replication fork stabilization/restart mechanisms, mainly MRE11 and EXO1 ^27,29^. MRE11 and EXO1 have been shown to be required for MUS81 and POLD3-mediated fork processing and restart ^28^. Interestingly, MUS81 is a key player in the complex mechanism termed “Fork cleavage-religation cycle” that has been shown to be required for fork restart specifically at R-loop-mediated stalled forks ^39^. Therefore, it is likely that DDK activity precedes the actions of MUS81 at R-loop-mediated stalled forks and is required for their proper restart. Ewing sarcoma cells have high CDK9 activity and are known to harbor large amounts of R-loops ^18^. It is likely that there is a high level of R-loop-mediated fork stalling. Therefore, the RS we are observing upon DDKi treatment in Ewing cells may be a result of a lack of fork restart at R-loop-mediated stalled forks. This also explains why CDK9 activity suppresses the cell cycle accumulation that occurs upon DDKi treatment in Ewing cells.

Previous studies looking at the effects of DDK inhibition on cell cycle have shown that TAK-931 treatment causes an increase in the level of p-HH3-positive cells in cancer cells ^30^; this is indicative of a DDK inhibitor-induced mitotic arrest. This result was not observed in Ewing sarcoma cells, supporting the idea that these cells may be able to enter and progress through mitosis with under-replicated DNA. Importantly, this phenomenon was independent of TP53 status, as the TC32 (WT TP53) cell line exhibited similar phenotypes when treated with DDKi (supplemental figure 7). A failure to complete replication should activate a G2/M checkpoint. While there was evidence of a G2/M accumulation upon DDKi treatment (**figure 3B & E right**), this does not seem to be a permanent arrest. Instead, Ewing cells seem to progress into and through mitosis. This results in the formation of abnormal mitotic structures including anaphase bridges, lagging chromosomes and the disorganization of centrosomes (**figure 5A & B**) likely due to an uncoupling of DNA replication and mitosis which has previously been proposed to be linked to a prolonged S-G2 phase ^30^.

Importantly, BRCA1 has been implicated in the enforcement of the G2/M checkpoint through the repression of cyclin B1 ^40^ and a recent report showed that EWS-FLI1 fusion-harboring cells are deficient for BRCA1 function due to increased retention of BRCA1 at active transcription complexes ^18^. It is, therefore, possible that Ewing sarcoma cells can enter mitosis prematurely in the face of under-replicated DNA due to RS. This also may explain the fact that there seems to be an accumulation of cells in the late-S and G2 phases of the cell cycle by 24 hours (**Figure 3B**) that is not accompanied by an accumulation of cells at the mitotic checkpoint, even at 48 hours of treatment. This may also underlie the sensitivity of Ewing sarcomas to chemotherapeutics that inhibit replication progression such as Gemcitabine and Aphidicolin ^17^ and opens the door to the development of novel therapeutic combinations that will greatly benefit the clinical outcomes of patients with Ewing sarcoma in the future.

## Materials and Methods

### Cell lines

The Ewing sarcoma cell lines that were used were A673, SK-ES-1 and RD-ES (ATCC). As a control non-Ewing sarcoma cancer cell line, the osteosarcoma cell line U2OS was used. A673 cells were cultured in Dulbecco’s modified Eagle’s medium (DMEM; Corning. Mediatech Inc.) supplemented with 10% fetal bovine serum (FBS) and 1% antibiotic-antimycotic (Gibco Life Technologies). U2OS cells were cultured in McCoy’s 5A (Corning. Mediatech Inc.) supplemented with 10% FBS and 1% antibiotic-antimycotic (Gibco Life Technologies). SK-ES-1 cells were cultured in McCoy’s 5A medium (Corning. Mediatech Inc.) supplemented with 15% FBS and 1% antibiotic-antimycotic (Gibco Life Technologies). RD-ES cells were cultured in RPMI-1640 (Corning. Mediatech Inc.) supplemented with 15% FBS and 1% antibiotic-antimycotic (Gibco Life Technologies). BJ cells were cultured in Minimum Essential Media (MEM; Corning. Mediatech Inc.) supplemented with 10% FBS and 1% antibiotic-antimycotic (Gibco Life Technologies). All cell lines were grown at 37°C and 5% CO_2_.

### Chemical Inibitors

The CDC7 kinase inhibitor, XL413, was purchased from Sigma-Aldrich (cat. No. SML1401; batch #: 0000036390) and dissolved in sterile filtered ddH_2_O to a final concentration of 5mM. The CDC7 kinase inhibitor TAK-931 (Simurosertib) was purchased from MedChemExpress (MCE) (cat. No. HY-100888) as a 10mM stock dissolved in DMSO.

### Cell viability

Cells were seeded at appropriate densities (adherent cell lines: 5,000 cells per well; mixed-adherent cell lines: 10,000 cells per well) and allowed to incubate overnight at 37°C. Cells were treated with serial dilutions of drug and incubated for 72 hours. Cell counting kit-8 (WST-8/CCK8) reagent from GLPBIO was added to each well and allowed to incubate for 2 hours. The absorbance was measured at 450nm on a SpectraMax M2/M2E microplate reader (Molecular Devices). Final cell viability was determined relative to the average absorbance of solvent control-treated cells.

### Western blotting

Cell lysates were separated by SDS-PAGE and then transferred to polyvinylidenedifluoride (PVDF) membranes. The membranes were blocked with 5% milk in TBS-T (TBS containing 0.1% Tween-20) for 1 hour at room temperature and then incubated in 1° antibody dilutions overnight at 4°C: rabbit anti-Cleaved PARP (Asp-214) #9451 (9541S), 1:1000; rabbit anti-MCM2 #4007 (4007S), 1:1000; Cell Signaling Technology, Inc; rabbit anti-GAPDH, ab9485; 1:5000; rabbit anti-gamma H2A.X (phospho S139), ab11174, 1:1000; rabbit anti-MCM2 (phospho S40+S41), ab70371, 1:1000; Abcam plc; rabbit anti-Phospho-Histone H3 (Ser10), Cell signaling technology Cat No.: 9701S, 1:1000. The following day, the membranes were washed 3 times for 10 minutes each in TBS-T and then incubated with 2° antibodies for 1 hour at room temperature: Goat anti-Rabbit IgG (H+L) Highly cross absorbed Secondary Antibody, Alexa Fluor Plus 800, Invitrogen #A32735, 1:40,000; Goat anti-Rabbit IgG (H+L) Highly cross absorbed Secondary Antibody, Alexa Fluor Plus 680, Invitrogen #A32734, 1:40,000; Goat anti-Mouse IgG (H+L) Highly cross absorbed Secondary Antibody, Alexa Fluor Plus 800, Invitrogen #A32730, 1:40,000; Goat anti-Mouse (H+L) Highly cross absorbed Secondary Antibody, Alexa Fluor Plus 680, Invitrogen #A32729, 1:40,000. Proteins were detected using an Odyssey western blot imaging system by LI-COR Biosciences. GAPDH was used as an internal loading control.

### Immunofluorescence microscopy

Cells were seeded onto EtOH-sterilized glass coverslips and allowed to attach for 24-48 hours. Following drug treatment, media was removed, and coverslips were washed in 1X PBS. For EdU incorporation assays, 10µM EdU was added to media 30 minutes prior to cell fixation. Cells were fixed with 4% paraformaldehyde for 15 minutes and permeabilized with 0.05% Triton-X100 for 15 minutes at room temperature. For EdU incorporation assays, click-It reaction was performed according to manufacturer’s instructions (Click-IT™ EdU Cell Proliferation Kit for Imaging, Alexa Fluor™ 488 dye, Cat. No. C10337). Coverslips when then blocked with 5% milk for 1 hour at room temperature and then incubated with 1° antibodies overnight at 4°C protected from light: rabbit anti-gamma H2A.X (phospho S139), ab11174, 1:500; rabbit anti-53BP1, ab36823, 1:500. Following the overnight incubation, coverslips were washed with 1X PBS for 10 minutes, 3 times and then incubated with 2° antibodies for 1 hour at room temperature protected from light: Goat Anti-Mouse IgG H&L (Alexa Fluor® 488) pre-absorbed, ab150117, 1:500; Goat Anti-Rabbit IgG H&L (Alexa Fluor® 488), ab150077, 1:500; Abcam plc. Goat anti-Mouse IgG (H+L) Cross-Absorbed Secondary Antibody, Alexa Fluor 568, A-11004, 1:500; Goat anti-Rabbit IgG (H+L) Cross-Absorbed Secondary Antibody, Alexa Fluor 568, A-11011, 1:500; ThermoFisher Scientific. The coverslips were then washed with PBS for 10 minutes, 3 times, stained with DAPI and mounted using ProLong™ Gold Antifade Mounting reagent (ThermoFisher Scientific, Cat no. P36930). Fluorescent images were captured on a Zeiss Axio Imager.A2 at 63x magnification. Corrected gamma-H2AX intensity and 53BP1 foci per cell for individual cells was measured using ImageJ software.

### Cell cycle

Cells were treated and then fixed in ice cold 70% ethanol at 4°C for 30 minutes. Cells were washed with 1X PBS and then incubated with Propidium Iodine/RNase solution for 2 hours. DNA content was then analyzed using flow cytometry. ModFit 5.0 software was used to analyze cell cycle based on DNA content.

### Annexin-V staining

Following drug exposure, cells were trypsinized, washed with PBS and resuspended in Annexin-binding buffer. Apoptotic cells were stained for annexin-V and DNA of dead cells was stained with propidium iodide according to the manufacturer’s instructions (Dead Cell Apoptosis Kit with Annexin V FITC and PI, for flow cytometry, ThermoFisher Scientific, Cat. No.: V13242). Cells were then analyzed using flow cytometry.

### Phospho-Histone H3 staining

Following treatment, cells were fixed in appropriate volume of 4% paraformaldehyde at room temperature for 15 minutes. Cells were then permeabilized by adding 100% ice-cold Methanol and incubating for 10 minutes on ice. Cells were then wash in PBS and resuspended in 100µL of 1° antibody (Phospho-Histone H3 (Ser10), Cell signaling technology Cat No.: 9701S, 1:50 dilution) for 1 hour at room temperature. Cells were then washed with PBS and resuspended in 100µL of diluted secondary antibody (Goat Anti-Mouse IgG H&L (Alexa Fluor® 488) pre-absorbed, ab150117, 1:500; Goat Anti-Rabbit IgG H&L (Alexa Fluor® 488), ab150077, 1:500; Abcam plc. Goat anti-Mouse IgG (H+L) Cross-Absorbed Secondary Antibody, Alexa Fluor 568, A-11004, 1:500; Goat anti-Rabbit IgG (H+L) Cross-Absorbed Secondary Antibody, Alexa Fluor 568, A-11011, 1:500; ThermoFisher Scientific). DNA content was labeled with propidium iodide as described in methods description of “cell cycle”. Cells were then analyzed using flow cytometry.

### Double-Thymidine Block

Cells were treated with 2mM Thymidine (Millipore Sigma, Cat. Num.: T9250) for 18 hours. Thymidine was then removed, and cells were washed several times with PBS. Media was replaced with Thymidine-free full serum media for 9 hours. Cells were then treated with 2mM Thymidine for an additional 18 hours at which point, Thymidine was washed out and media was replaced +/- specified inhibitors for varying amounts of time.

## Supplemental Figure legends

**Supplemental figure S1.**
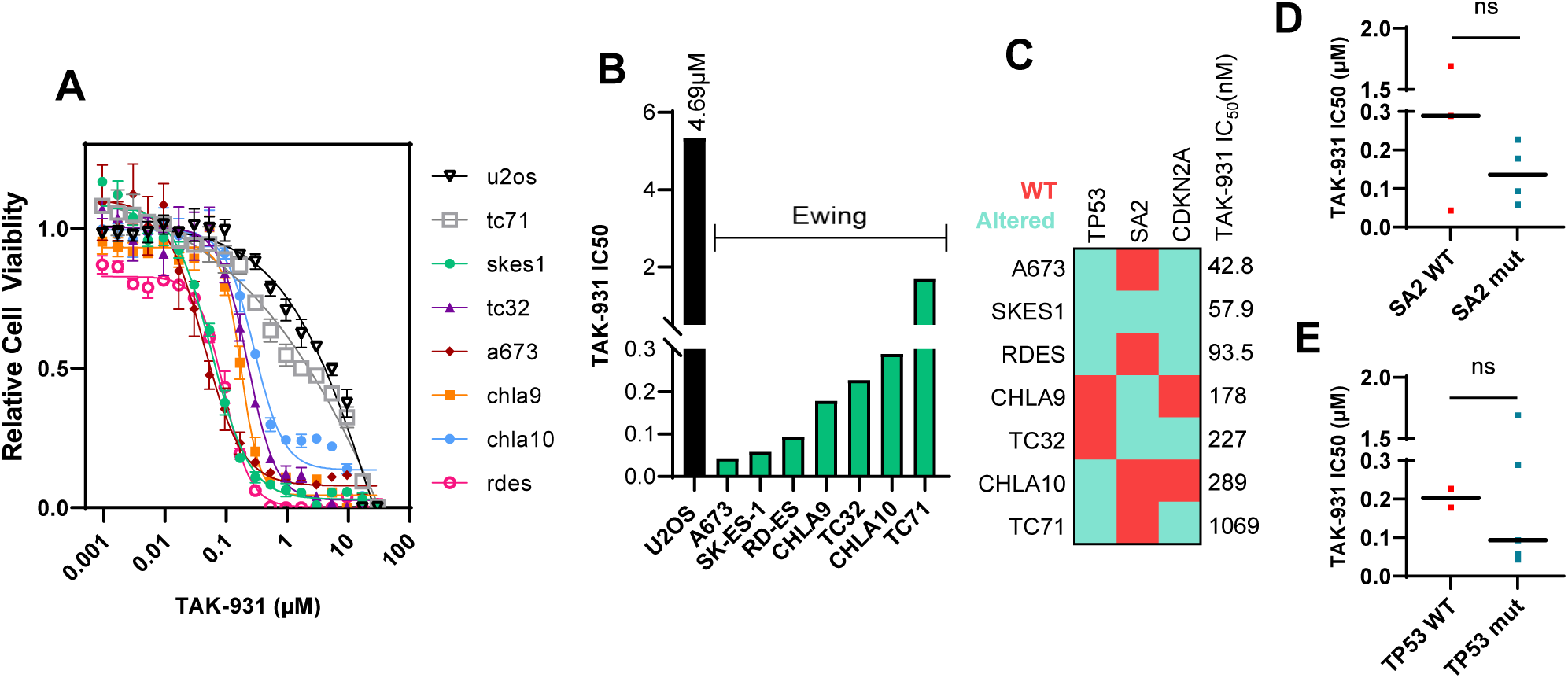
Ewing sarcoma cells are sensitive to the DDK inhibitor TAK-931 intendent of TP53. STAG2 and CDKN2A status. A) A panel of Ewing sarcoma cell lines (colored lines) and the osteosarcoma cell line U2OS (black line) were treated with a wide range of doses of TAK-931 for 72 hours. Cell viability was assessed using CCK8 reagent and relative viability was calculated based on DMSO treated control cells. B) IC50 values were calculated using GraphPad prism software (non-linear regression [Inhibitor] vs. Response – Variable slope (four parameters)) for TAK-931 based on relative viability curves generated in panel A. Green bars represent all Ewing sarcoma cell lines and black bar is U2OS non-Ewing control. C, D & E) IC50 values were compared between cell lines of different genetic backgrounds (TP53, STAG2 and CDKN2A status). C) Cell lines were arranged from based on TAK-931 sensitivity from most sensitive (top) to least sensitive (bottom). A colored grid was then generated using red squares to represent WT gene status and blue squares to represent altered (mutant/deleted) gene status. No apparent trends were observed. D) Red dots represent IC50 values of cells with WT STAG2 and blue dots represent IC50 values of cells with mutant STAG2. E) Red dots represent IC50 values of cells with WT TP53 and blue dots represent dots IC50 values of cells with mutant TP53.

**Supplemental Figure S2.**
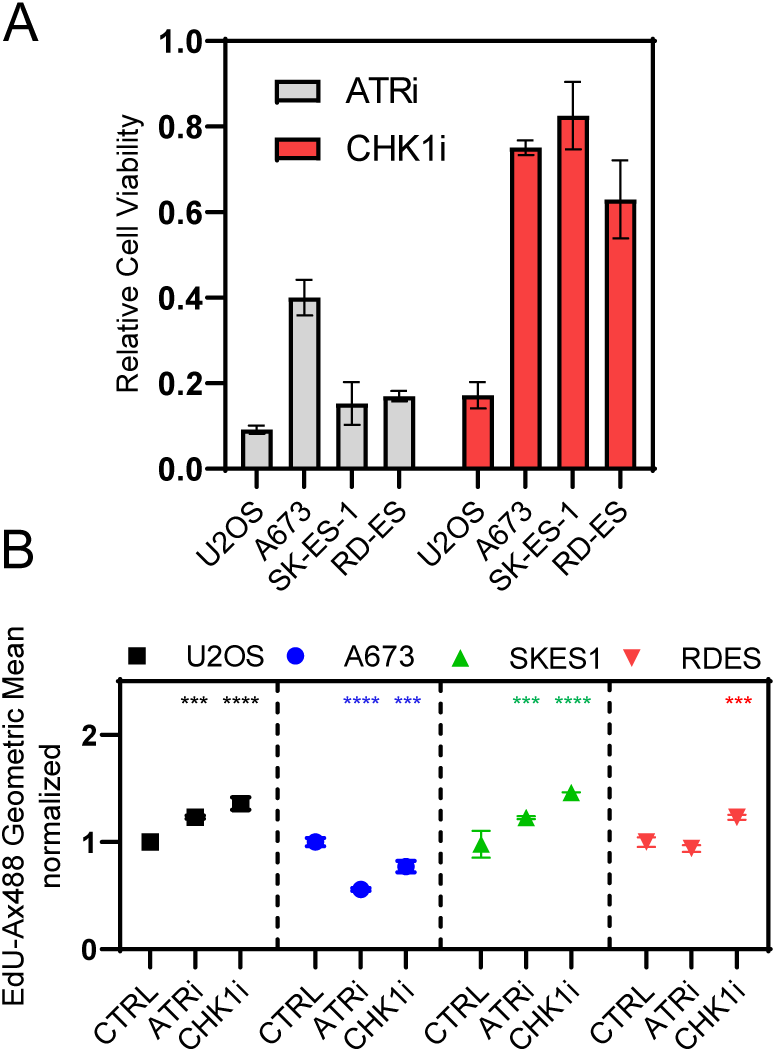
Ewing Sarcoma cells are not sensitive to ATR or CHK1 inhibitors as single agents. A) Ewing Sarcoma cells and U2OS cells were treated with the 5µM of the ATR inhibitor AZD6738 or 1µM of the CHK1 inhibitor LY2603618 for 72 hours and cell viability was assessed using the CCK8 cell counting kit-8 reagent. Relative cell viability was calculated based on DMSO treated cells (n = 3 technical replicates). B) Ewing Sarcoma cells and U2OS cells were treated with 5µM AZD6738 or 1µM LY2603618 for 2 hours followed by 10µM EdU for 30 minutes. Cells were fixed, stained and EdU incorporation rate was measured using flow cytometry. (n = 2 biological replicates.) (***p<0.001, ****p<0.0001).

**Supplemental Figure S3.**
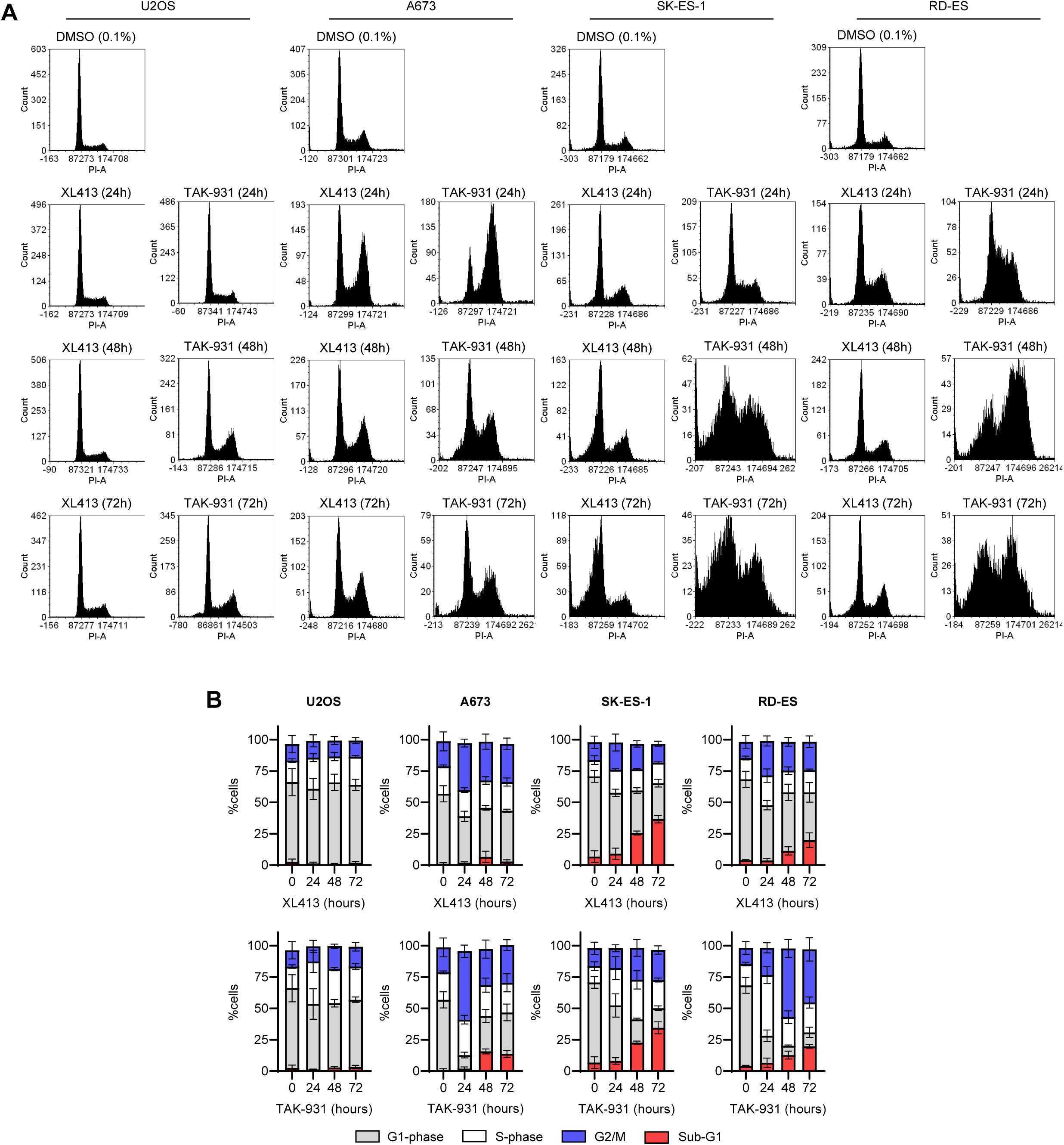
DDK inhibition causes aberrant cell cycle accumulation in Ewing Sarcoma cells. Ewing Sarcoma cells and U2OS cells were treated with 0.1% DMSO (72 hours), 300nM TAK-931 or 1µM XL413 for 24, 48 and 72 hours, DNA was stained with propidium iodide and DNA content was analyzed using flow cytometry. Cell cycle distribution was measured using FCS Express v7. A) Representative histograms of three independent experiments B) Quantification of cell cycle distribution; G1 = grey bars, S = white bars, G2/M = blue bars, sub-G1 = red bars (n=3 biological replicates).

**Supplemental Figure S4.**
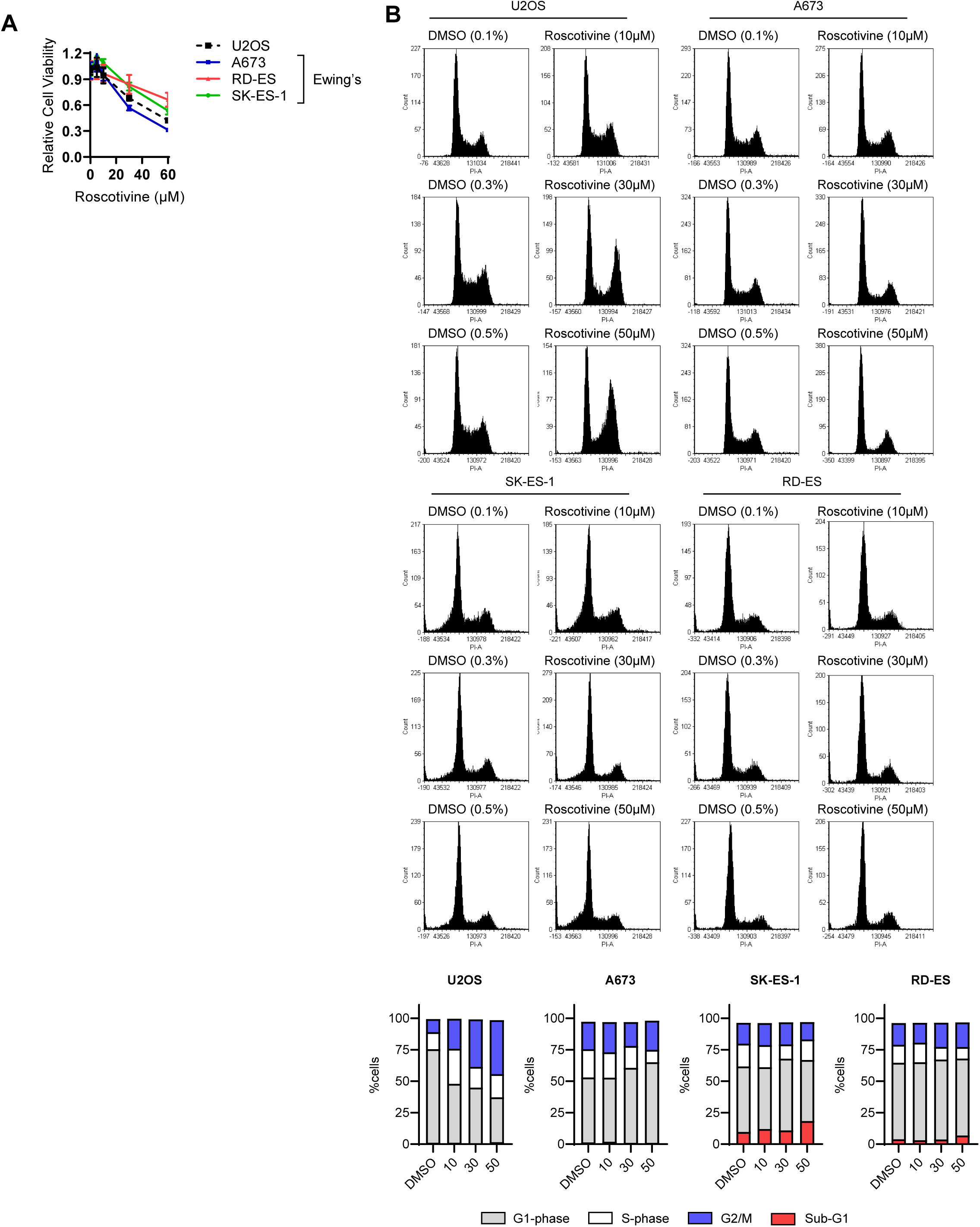
DDKi-induced affects are suppressed by inhibition of CDK9. A &B) Ewing sarcoma cells were treated with 300nM TAK-931 or 1µM XL413 +/- 20µM DRB for 24 hours. DNA content was analyzed using flow cytometry. A) Representative histogram images of A673 cells of 2 independent experiments. B) Cell cycle quantification of A673 (top), RD-ES (middle) and SK-ES-1 (bottom) under the specified conditions (n=2 biological replicates).

**Supplemental Figure S5.**
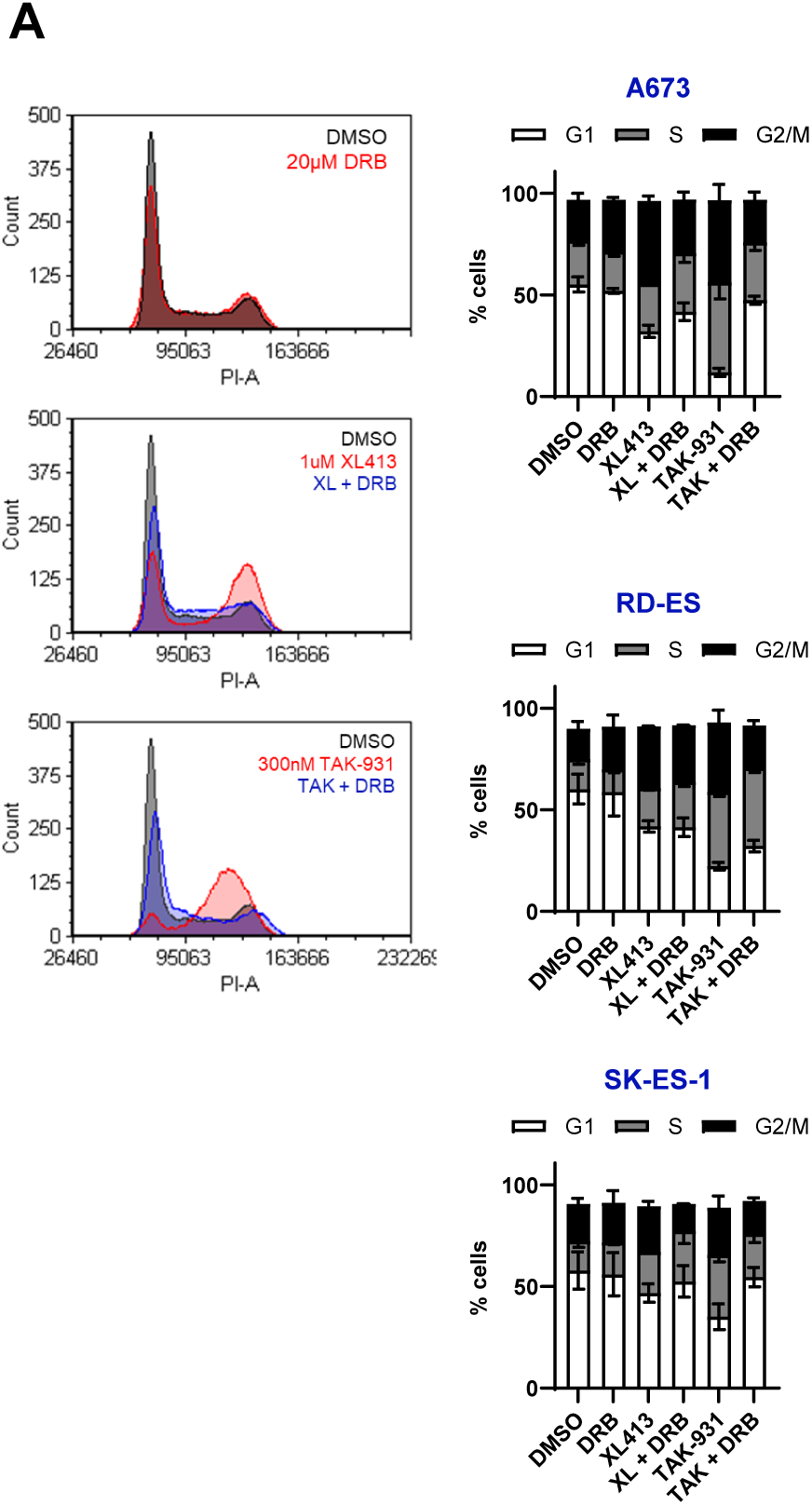
Ewing Sarcoma cells are not sensitive to the CDK2 inhibitor Roscotivine. A) Ewing Sarcoma cells and U2OS cells were treated with increasing concentrations of the CDK2 inhibitor Roscotivine for 72 hours and cell viability was measured using CCK-8 cell viability reagent. Relative viability was calculated based on DMSO treated cells. (n=3 technical replicates). B) Ewing Sarcoma cells and U2OS cells were treated with increasing concentrations of DMSO and Roscotivine for 24 hours, DNA was stained with propidium iodide and DNA content was analyzed using flow cytometry. Cell cycle distribution was measured using FCS Express v7. Quantification of cell cycle distribution; G1 = grey bars, S = white bars, G2/M = blue bars, sub-G1 = red bars (n=3 biological replicates).

**Supplemental Figure S6.**
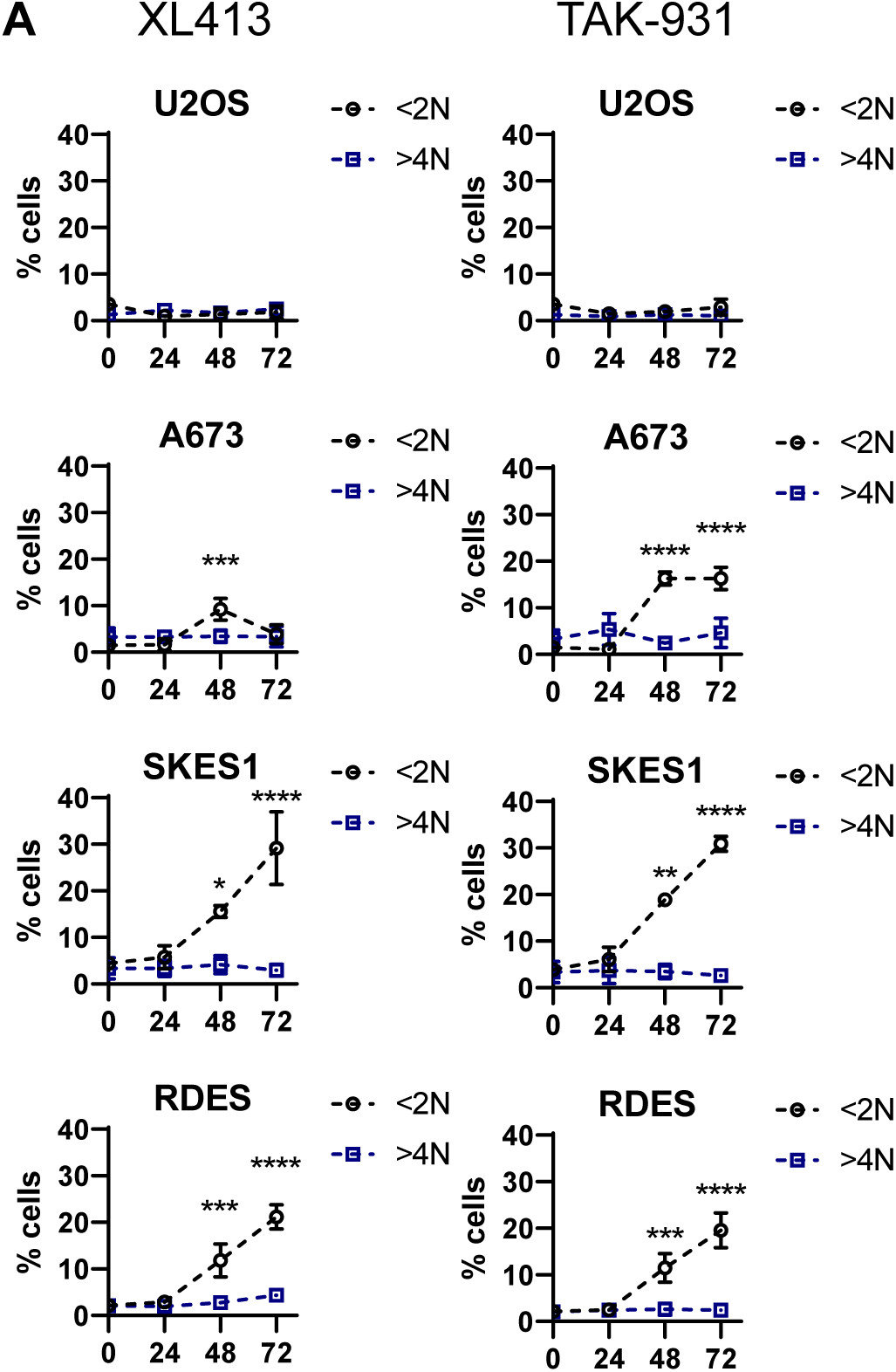
DDK inhibition does not result in significant aneuploidy in Ewing Sarcoma cells. Ewing Sarcoma cells and U2OS cells were treated with 0.1% DMSO (72 hours), 300nM TAK-931 or 1µM XL413 for 24, 48 and 72 hours, DNA was stained with propidium iodide and DNA content was analyzed using flow cytometry. Percentages of cells with <2N (black dotted line) and >4N (blue dotted line) was measured using FSC Express v7 (n=3 biological replicates) Ordinary one-way ANOVA, *p<0.05, ***p<0.001, ****p<0.0001.

**Supplemental Figure S7.**
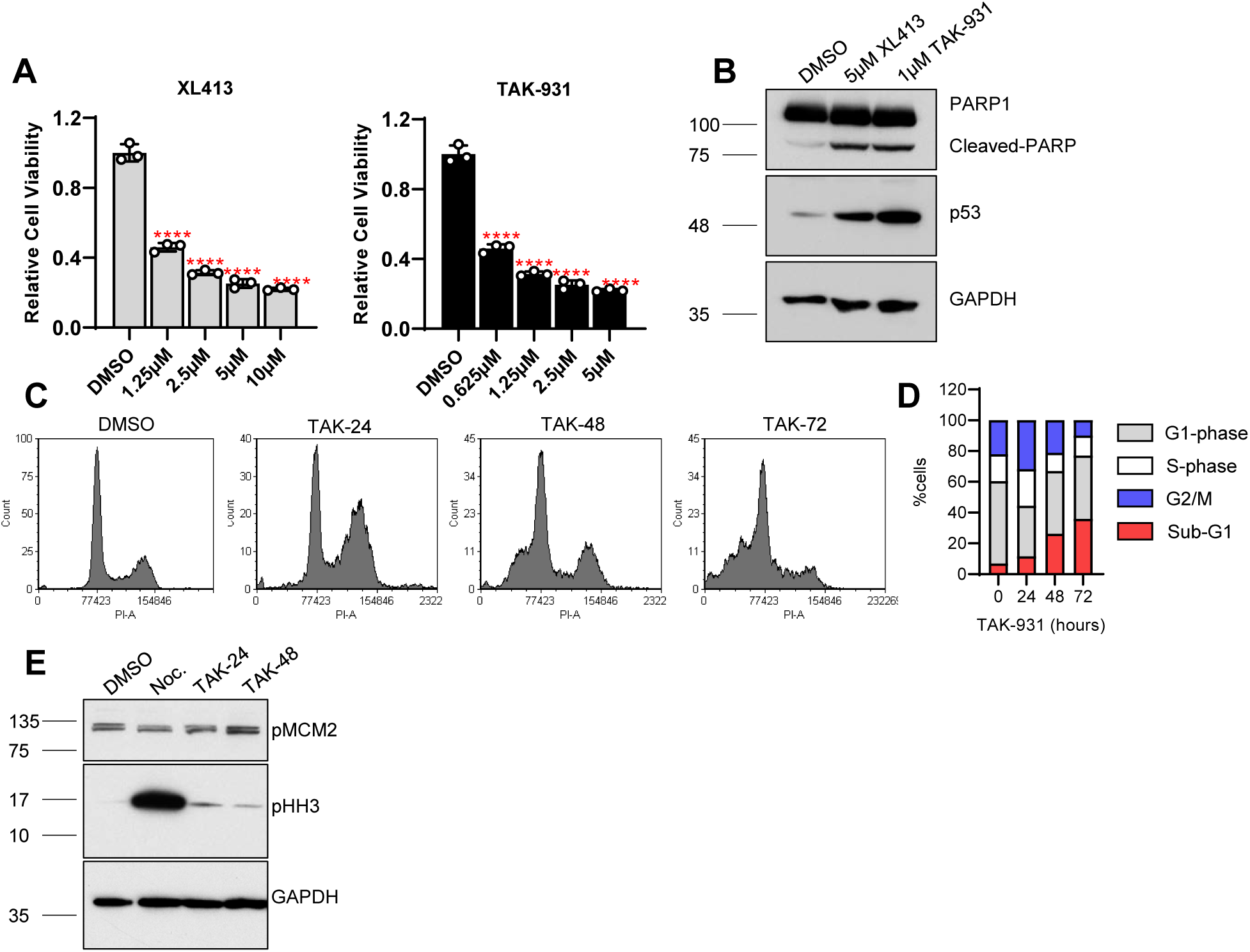
TP53-WT TC32 Ewing sarcoma cells are sensitive to DDKi. A) TC32 cells were treated with specified concentrations of DDKi - XL413 (left-grey bars) and TAK-931 (right – black bars) for 72 hours and cell viability was assessed using CCK8. Relative viability was calculated based on DMSO treated controls (n = 3 technical triplicates; Ordinary one-way ANOVA compared to DMSO treated control ****p<0.0001). B) TC32 cells were treated with either 0.1% DMSO, 5µM XL413 or 1µM TAK-931 for 48 hours. Protein lysate was collected, and a western blot was run for specific proteins. GAPDH was used as a loading control (n = 1). C) TC32 cells were treated with 300nM TAK-931 for specified times (DMSO = 0.1% DMSO for 72 hours). DNA content histograms were generated using flow cytometry and cell cycle distributions were quantified in panel D. E) TC32 cells were treated with either 0.1% DMSO, 1µM Nocodazole (24 hours) or 300nM TAK931 for 24 or 48 hours. Protein lysate was collected, and a western blot was run for specific proteins. GAPDH was used as a loading control (n = 1). Upon DDK inhibition, TC32 cells (TP53 WT) experience a similar reduction in cell viability, pattern of cell cycle delay and apoptosis asTP53 mutant cells. Importantly, these cells, too, do not experience a mitotic accumulation, suggesting that the lack of mitotic delay upon DDK inhibition in A673 cells is not due to the lack of p53 function.

## Author Contribution Statement

JO supervised the work and revised the manuscript. JM designed and performed experiments, analyzed and interpreted data, provided statistical analysis and contributed the bulk of the manuscript text. AG and TH provided important clinical insight and assisted in the drafting and revision process. TM, AW, JK, JK and LG provided crucial molecular insight and assisted in the drafting and revision process. ML provided statistical expertise and assisted in the drafting and revision process. All authors reviewed the final version of the manuscript and approve of its contents.

## Ethics Statement

This study did not require ethical approval.

